# The evolutionary lifecycle of ribosome hibernation factors

**DOI:** 10.1101/2025.11.04.686544

**Authors:** Lewis I. Chan, Kimiho Omae, Charlotte Rebecca Brown, Kazuaki Amikura, Chinenye L. Ekemezie, Jessica Clark, Kenji Nishide, Karla Helena-Bueno, Pawel Palmowski, Andrew Porter, Shino Suzuki, Sergey V. Melnikov

**Author notes:** Contributed equally (Lewis I. Chan, Kimiho Omae, Charlotte Rebecca Brown, Kazuaki Amikura).

## Abstract

Bacteria defend against hostile environments through a variety of molecular mechanisms, including ribosome hibernation. Previously, bacteria were shown to initiate ribosome hibernation by activating protective proteins known as hibernation factors. It was demonstrated that hibernation factors prevent ribosome degradation by nucleases, which allows bacteria to safely store their inactive ribosomes and survive under starvation or persistent stress. Because homologs of hibernation factors were found in diverse lineages of bacteria, it is currently assumed that the mechanism of ribosome hibernation is highly conserved across species. Here, we assess 46,015 complete bacterial genomes to reveal the principles underlying the origin and evolution of these essential proteins in bacterial cells. We find that hibernation factors emerged in ancient bacteria as relatively large proteins that then gradually reduced in size and have undergone complete extinction in over 10% of studied bacteria. We then demonstrate that the degeneration of ancient hibernation factors is often accompanied by “borrowing” hibernation factors from other species via gene transfers, *de novo* gene birth or fusion of truncated hibernation factors with fragments from other stress-response proteins. These findings reveal a unique evolutionary pathway in which bacteria respond to the reductive evolution of hibernation machinery by inventing novel hibernation mechanisms, thus restoring their capacity to survive starvation and stress. This model implies that most ribosome hibernation factors are yet to be discovered and predicts the organisms that rely on currently unknown hibernation mechanisms.

## INTRODUCTION

The synthesis of ribosomes represents the most metabolically expensive event in the cell as actively growing bacteria invest up to 60% of their energy into the production of these essential macromolecules.^1,2^ It is therefore not surprising that living cells of diverse species have evolved a specialized mechanism, known as “ribosome hibernation”, which allows cells to safely store their ribosomes under conditions of stress and starvation—so that most other cellular macromolecules degrade but ribosomes remain intact for long periods of times that can exceed decades.^3^

Previous studies revealed that bacteria initiate ribosome hibernation by activating proteins known as hibernation factors.^3–15^ Under conditions of stress or starvation—when cells inactivate protein synthesis and accumulate idle ribosomes—hibernation factors bind to the active sites of ribosomes and prevent ribosome degradation by nucleases. As a result, starved or persistently stressed bacteria can endure prolonged metabolic inactivity with a lower risk of death due to irreparable damage to essential cellular machinery.

The critical role of hibernation factors in bacterial survival has been previously demonstrated in a handful of model organisms, primarily *Escherichia coli*, which was shown to bear seven hibernation factor proteins (HPF, RaiA, RMF, YqjD, ElaB, Sra, and YgaM). The importance of these factors is evident from their abundance during dormancy: in stationary phase, mRNAs encoding hibernation factors constitute up to 25% of the total transcriptome^16^, and these proteins themselves account for up to 7% of the total cellular protein mass.^17^ This places them among the most abundant proteins in non-growing cells, with their intracellular concentration exceeding that of ribosomes by an order of magnitude.^18^ Genetic deletion studies showed that the loss of even a single hibernation factor, such as the first-to-be-characterized factor RMF, reduces cell viability by up to 1,000-fold in stationary phase or under persistent environmental stress (e.g., cold or acid shock).^19,20^ This fitness defect is compounded several orders of magnitude further when multiple factors are deleted simultaneously.^21^ Thus, ribosome hibernation factors are indispensable for sustainable bacterial survival in hostile environments and the continuity of microbial life on Earth.

Until recently, mechanisms of ribosome hibernation in bacteria were viewed as largely conserved and inevitably relying on the ubiquitous hibernation factors from the HPF/RaiA family (also known as HPF, long-HPF, RaiA, RafH, mPy and others).^22– 28,18^ However, more recently, alternative ribosome hibernation mechanisms were discovered, involving lineage-specific hibernation factors RTOF^29^ and Balon^14^, as well as archaeal hibernation factors aRDF, Dri and Hib, which were acquired by certain bacteria via horizontal gene transfers.^30–32^ These studies revealed that ribosome hibernation in bacteria can be mechanistically diverse between species, but despite the years of extensive studies we currently do not know: Why do some bacteria have alternative ribosome hibernation mechanisms? What species are most likely to possess currently unknown ribosome hibernation strategies? When did currently established hibernation factors emerge in bacterial species? And what principles guide the evolution and spread of these essential proteins across species?

In this study, we provide answers by combining evolutionary, proteomic, and genetic analyses to trace the origin and evolutionary journey of ribosome hibernation factors throughout 4.3 billion years of bacterial existence. Using homology searches combined with phylogenetic dating, we first determine the absolute age of each family of ribosome hibernation factors found in modern *E. coli*, illustrating their gradual emergence over a span of 3.9 billion years. We then reconstruct the ancestral forms of ribosome hibernation factors to infer when these factors emerged in ancient bacteria. Our analysis reveals that most bacteria tend to evolve ribosome hibernation factors as larger proteins that undergo gradual degeneration, accumulating truncations and, in over 10% of species, even complete protein loss over time. We further find that this degeneration appears to force organisms to reinvent and diversify their ribosome hibernation machinery— either by evolving new hibernation factors or by acquiring their genes from other species via horizontal gene transfers, explaining the origin of the high degree of redundancy of the ribosome hibernation machinery observed today. These findings reveal a unique evolutionary pathway in which bacteria respond to the natural degradation of their hibernation machinery by inventing novel hibernation mechanisms, thus restoring their capacity to survive starvation and stress. This model implies that most key mechanisms of ribosome hibernation are yet to be discovered and predicts organisms that are most likely to possess currently unknown hibernation factors.

## RESULTS AND DISCUSION

### Most ribosome hibernation factors are young proteins

We first set out to determine the age of ribosome hibernation factors and the order in which these protective proteins emerged in bacteria. To approach this, we analyzed 46,015 complete bacterial genomes to identify all detectable homologs of hibernation factors initially discovered in *E. coli* and then reconstructed their evolution by tracing similarities of their sequences, structures, as well as their corresponding operons in representative species (**Data S1, Materials and Methods**). Our analysis was driven by the fact that all previous studies were based on merely mapping detectable homologs of hibernation factors on the tree of life without separating their strictly vertically evolving homologs from those evolving via horizontal gene transfers^14,29,30,33–35^. As a result, it was impossible to answer when hibernation factors emerged in bacteria, how these factors spread across bacterial genomes, or why certain bacteria possess dissimilar sets of hibernation factors compared to their close relatives on the tree of life.

To resolve this, we first separated homologous hibernation factors into pools of vertically evolving proteins. For example, rather than grouping proteins HPF and RaiA (in *E. coli*) into a single family of HPF/RaiA proteins—as commonly practiced in the field^14,29,30,33–35^—we have separately traced the evolution for the HPF lineage and the RaiA lineage by assessing the conservation of protein sequences and gene operons (**Fig. 1a, Figs. S1-4**). We then completed similar analyses for other hibernation factors, including YqjD, ElaB, YgaM, RMF and Sra, which allowed us to identify when each hibernation factor emerged in ancestral bacteria (**Fig. 1a,b**).

**Figure 1.**
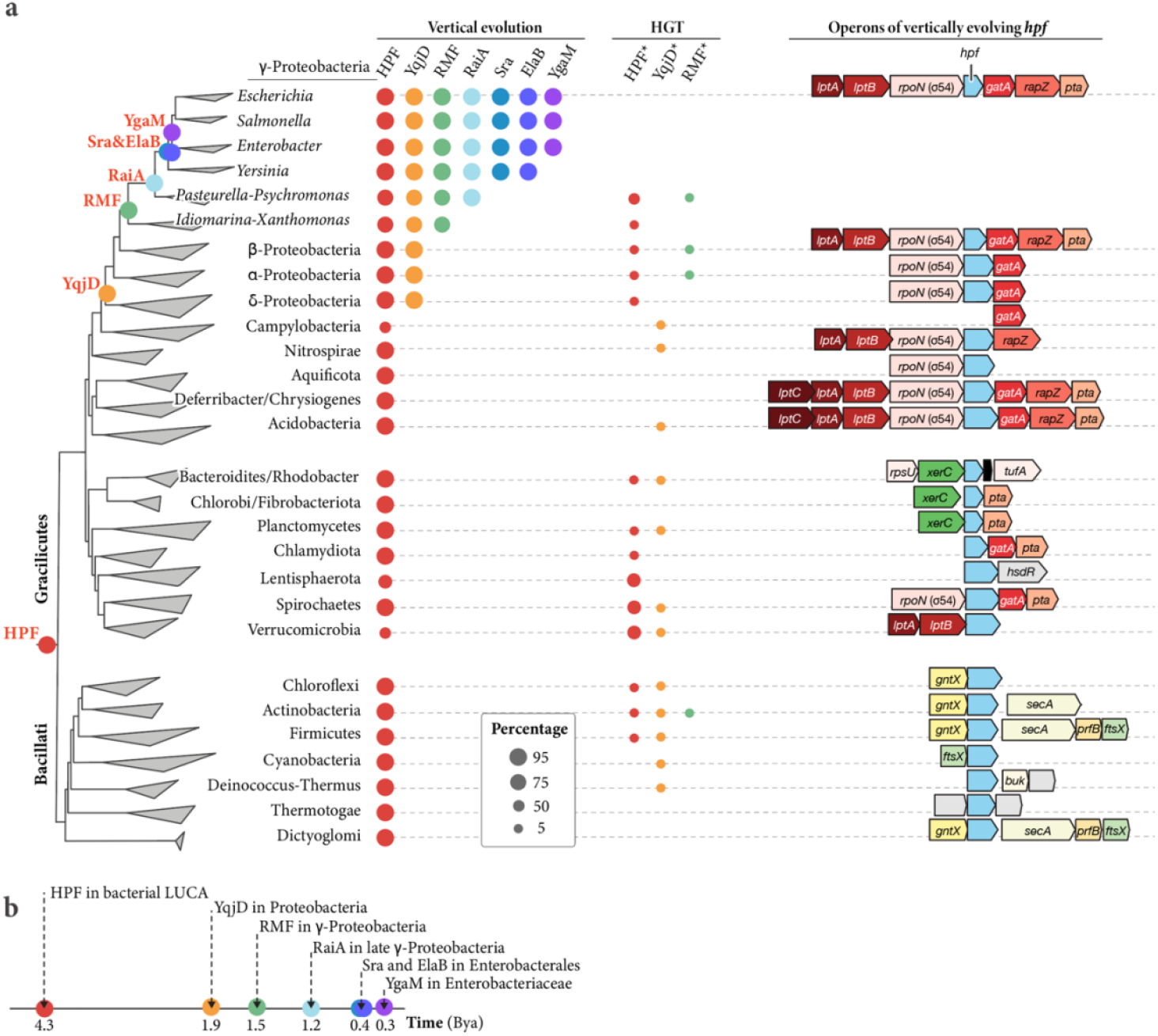
Most ribosomes hibernation factors are evolutionary young proteins. (**a**) The phylogenetic tree indicates apparent time points at which the ancestors of modern *E. coli* have acquired each of the ribosome hibernation factors. The right side of the panel shows the most typical operon structure for the ubiquitous hibernation factors HPF. (**b**) A schematic timeline illustrating the order in which ribosome hibernation factors have emerged, indicating that modern *E. coli* have acquired their hibernation machinery over a period of 3.9 billion years. Previous studies based on the domain occurrence, without reference. Although informative for the occurrence, they were not good enough to determine the age and predict the occurrence of these factors in a given lineage of bacteria. Overall, the figure illustrates that most ribosome hibernation factors of modern *E. coli* have originated relatively recently, after the separation of the bacterial phyla.

We found that most ribosome hibernation factors have emerged relatively recently in bacterial cells, and their age tends to correlate with their size, with more recently emerged factors being smaller in size (**Fig. 1a,b**). The protein identified in *E. coli* as HPF represents a direct descendant of the oldest hibernation factor in bacteria (which we refer to as ancient HPF). We could detect vertically evolving homologs of ancient HPF in most species of each phylum, pointing to its emergence in the last common ancestor of bacteria, approximately 4.3 billion years ago (Bya)^36^. This finding showed that bacteria acquired their ability to preserve ribosomes through hibernation at the very dawn of bacterial evolution. However, the remaining six hibernation factors found in modern *E. coli* have emerged much more recently, only after the origin of Proteobacteria: 2.4 billion years after the origin of ancient HPF, when the common ancestor of Proteobacteria acquired the YqjD gene. This was followed by the origin of the RMF gene, which occurred shortly after the emergence of the order Xanthomonadales, approximately 1.5 Bya. And only after this event, approximately 1.2 Bya, the ancestor of species spanning orders Enterobacteriales to Alteromonadales acquired the gene for RaiA, either via horizontal gene transfer from outside Proteobacteria or an ancient event of gene duplication (**Fig. S1-S3**). The most recently acquired hibernation factors, Sra (of likely *de novo* birth), ElaB (gene duplication) and YgaM (gene duplication), arose within Enterobacteriaceae, very recently, between 0.3 and 0.4 Bya.

Our analysis revealed that ribosome hibernation factors are substantially younger than currently thought. For instance, we showed that the previously observed presence of YqjD/ElaB/YgaM proteins across a broad range of phyla is not a result of their origin in the common ancestor of bacteria^35^, but of recent horizontal gene transfers from Proteobacteria (**Fig. S4**,**S5**). Similarly, protein RaiA considered specific to γ-proteobacteria^33,34^, have emerged substantially more recently, only after the emergence of Idiomarina branch, meaning that it is absent in about half of known γ-proteobacteria. These relatively recent origins shows that most ribosome hibernation mechanisms are phyla-or even sub-phylum specific rather than broadly conserved, implying that other phyla (e.g. Actinobacteria or Bacteroidetes) may currently have unappreciated strategies for ribosome hibernation.

### “Younger” hibernation factors tend to have a higher impact on bacterial starvation tolerance

Having found that the age of hibernation factors in a single organism can differ by as much as 4 billion years, we then asked: does the protective activity of hibernation factors depend on their evolutionary age? In other words, do evolutionarily ancient and ubiquitous hibernation factors play a more fundamental role in maintaining cellular dormancy, while younger, phyla-specific factors simply fine-tune this process?

To answer this, we compared the rate of *E. coli* recovery from stationary phase between the wild-type strain and strains with genomic deletions of individual ribosome hibernation factors (**Fig. 2a,b, Data S2**). We found that deletion of the most ancient hibernation factor, HPF, caused a detectable defect in growth recovery: after five days in stationary phase, HPF-less cells required 1 hour longer to show visible signs of growth compared to the wild-type strain. However, deletion of the genes for younger hibernation factors showed even stronger recovery defect, including: ∼1.5 hours delay for YqjD (1.9 Bya), ∼2 hours delay for RaiA (1.5 Bya), and 4 hours delay for RMF (1.8 Bya). Thus, we found that hibernation factors that are younger (and limited to an individual phylum) tend to have a comparable if not stronger impact on bacterial viability in stationary phase compared to the ancient HPF—illustrating that phylum-specific mechanisms of ribosome hibernation play the primary role in maintaining the burden of ribosome hibernation in bacterial cells.

**Figure 2.**
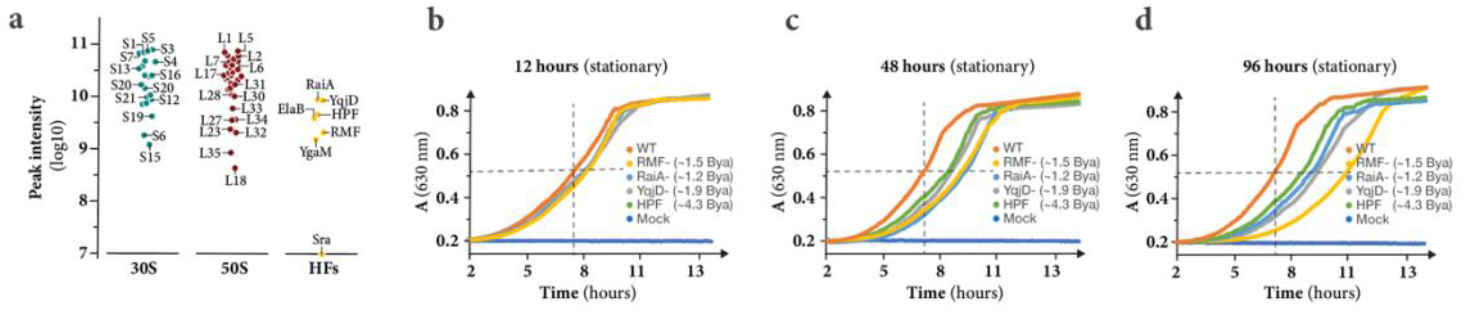
Evolutionary “younger” hibernation factors tend to have a higher fitness cost in dormancy. (**a**) Proteomic analysis of ribosomes isolated from stationary bacteria (∼24 hours in stationary phase) demonstrates the relative abundance of each ribosome hibernation factor (labelled as “HFs”) on the ribosome compared to the proteins of the large and small ribosomal subunits (labelled as “50S” and “30S”, respectively). (**b-d**) Growth curves illustrate the impact of genetic knockouts of individual ribosome hibernation factors on the rates of *E. coli* recovery from stationary phase of three different durations: 12 hours (**b**), 48 hours (**c**) and 96 hours (**d**). The crossing dashed lines indicate the point when the wild-type strain reaches the highest rate of division. The panel illustrates that both the wild-type *E. coli* and mutants lacking genes for hibernation factors (one factor at a time) recover progressively slower as the stationary phase gets longer, with deletions of “younger” hibernation factors having a comparable and typically stronger defect of growth recovery compared to the ancient hibernation factor HPF.

### Ribosome hibernation factors undergo accelerated evolution and degeneration

Because even the youngest hibernation factors emerged hundreds of millions of years ago, we next asked: how did the structure of ancient hibernation factors compare to those found in modern bacteria? Our motivation to answer this question arose from previous studies of protein HPF, showing that it exists in two isoforms—long and short—that initiate mechanistically dissimilar forms of ribosome hibernation. In species like *Thermus thermophilus, Bacillus subtilis, Staphylococcus aureus*, and *Lactobacillus lactis*, HPF was found as a long isoform (∼200 amino acids), which triggers the formation of nuclease-protected ribosome dimers, whereas in organisms like *E. coli* HPF was found as a short isoform (95 amino acids), which can form ribosome dimers and initiate efficient ribosome storage only with the help of an auxiliary hibernation factor, RMF.^37^ Therefore, we wanted to understand whether HPF emerged as a smaller protein that later acquired an additional domain, or as a longer protein that underwent truncations in some species, including *E. coli*.

To answer this, we assessed how the size of hibernation factors depends on the age of bacterial branches on the tree of life. We first found that the occurrence of short and long protein isoforms is highly atypical for canonical factors of protein synthesis (e.g. EF-Tu or EF-G), but is common not only for HPF, but for all other hibernation factors (**Fig. 3a,b**). Furthermore, we found that longer isoforms are common for more ancient and earlier-branching clades of species, indicating that hibernation factors have likely emerged as larger proteins that underwent gradual reduction in size in some of the recently emerging bacterial branches. For example, in Proteobacteria, the ancestral hibernation factor HPF has undergone a truncation of its C-terminal domain in the common ancestor of *β*- and *γ*-proteobacteria. Subsequently, in *γ*-proteobacteria, HPF experienced a gradual reduction in size upon transition from approximately 120 amino acids in early-branching proteobacteria (e.g. Xanthomonadales) to about 95 amino acids in more recently diverged lineages (e.g. Enterobacteriales) (**Fig. 3c**). Aside from Proteobacteria, we observed that HPF has acquired the truncation of the C-terminal domain in about a third of currently characterized bacterial phyla (**Fig. 3c**). Similar reductive evolution has occurred in other hibernation factors, including Sra, RMF and YqjD: these factors are prone to losing their N- and C-termini, with C-terminal protein segments being lost most frequently in each of them (**Fig. S6**). Thus, we found that hibernation factors tend to emerge as larger proteins that undergo gradual truncations in a substantial number of bacterial species.

**Figure 3.**
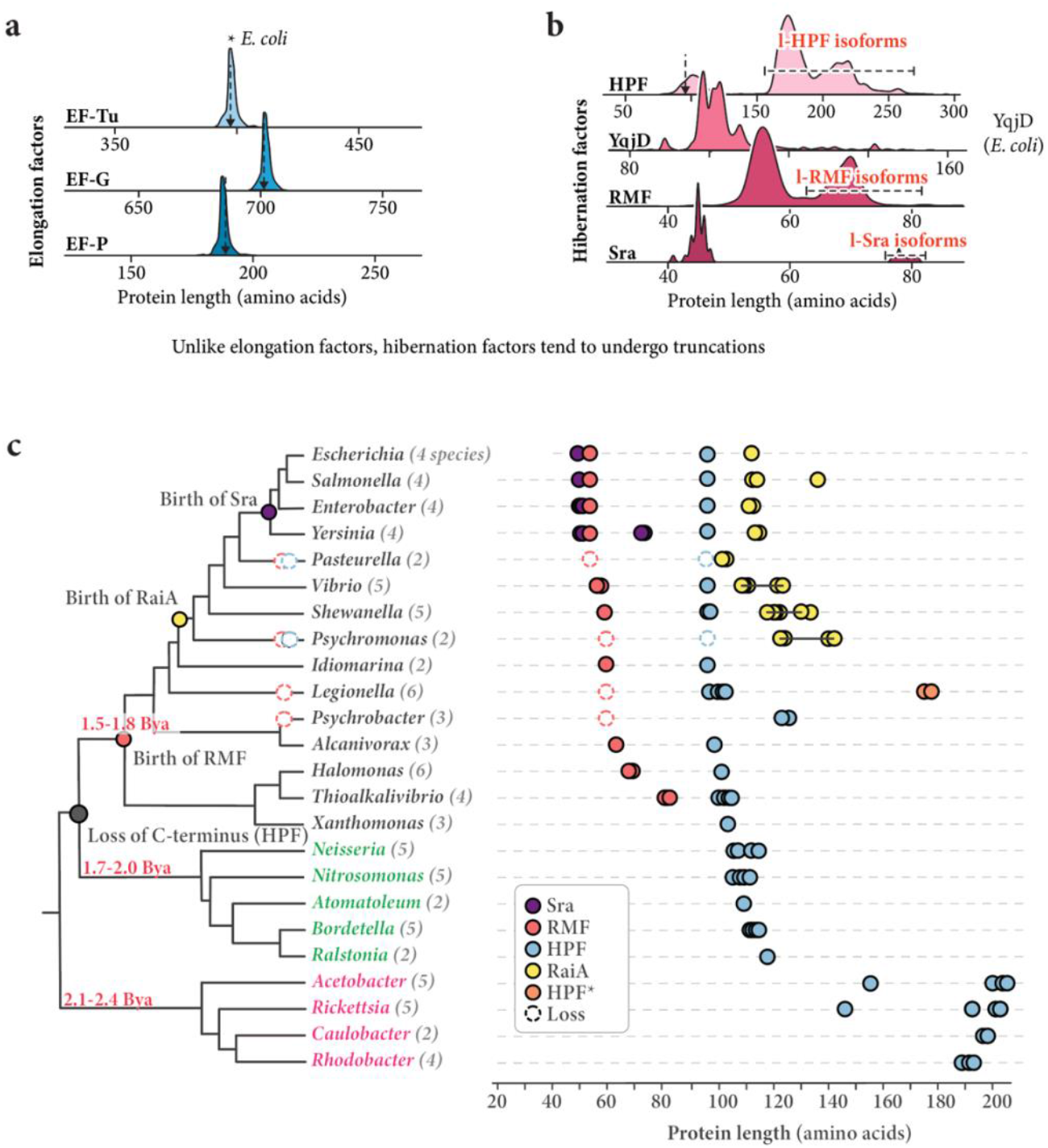
Bacteria tend to gradually degenerate their hibernation factors due to protein truncations or complete gene loss. (**a, b**) Plots illustrating the distributions of elongation factors (**a**) and hibernation factors (**b**) by their amino acid lengths across bacterial species. The panel shows that—by contrast to elongation factors— hibernation factors tend to exist in two forms: the longer forms (which typically correspond to more ancient protein variants), and truncated forms (that are occasionally found in bacterial lineages across the bacterial tree of life). (**c**) The tree illustrating that longer ribosome hibernation factors tend to undergo progressive reduction in size upon transition from earlier branching to later branching bacterial clades.

To understand possible evolutionary forces driving these truncations, we analyzed the ratios between non-synonymous and synonymous substitutions in the genes for ribosome hibernation factors to determine selective pressures acting on them. We reasoned that—if hibernation factors are only active in non-growing cells but remain functionally silent in actively growing cells—then mutations in their genes, even deleterious ones, should remain neutral for cellular fitness for as long as bacterial cells continue to grow and avoid dormancy. We therefore anticipated that actively growing cells may accumulate deleterious mutations in hibernation factor genes during active cell division, making hibernation factors evolve faster and under lower selective pressure compared to proteins involved in active cellular growth.

Our analysis revealed that ribosome hibernation factors indeed have sequences evolving at least eight times faster and under twice as low selective pressure compared to factors of protein synthesis —as evidenced by the overall rates of amino acid substitutions and the ratio between synonymous and non-synonymous mutations (**Fig. S7**). For example, the elongation factors EF-Tu and EF-G from closely related species *E. coli* and *Klebsiella pasteurii* have substitutions in only 1.5% and 3.8% of amino acids (96.2-98.5% of sequence identity) and the dN/dS ratios of 0.076 and 0.102— indicative of both high conservation and high purifying selective pressure. However, the hibernation factor HPF has substitutions in 12.6% of amino acids and the dN/dS ratio of 0.174, indicative more than eight times higher rates of evolution and a two times weaker selective pressure. Furthermore, factors YqjD and Sra display even higher rates of sequence evolution, exhibiting substitutions in 24% and 44% of their sequences between *Klebsiella pasteurii* and *E. coli* (compared to 1.5% for EF-Tu homologs).

Thus, we found that frequently occurring truncations of ribosome hibernation factors appear to reflect the natural and previously unappreciated cellular tendency to gradually degenerate these protective proteins, possibly due to their inactivity in actively growing (i.e. evolving) cells.

### Ancestral hibernation factors have undergone extinction in a substantial fraction of bacteria

Since we found that hibernation factors undergo progressive truncation in protein structure, we next asked whether this reductive evolution can result in their complete loss in some bacterial species. Our analysis of bacterial genomes revealed that this sporadic gene loss is widespread across bacteria (**Fig. 4a, Data S3**). For instance, even among the closest relatives of *E. coli* (Enterobacterales)—where the *hpf* and *rmf* operons are conserved in most species—the *hpf* and *rmf* genes are absent in at least 25 distinct genera, including species of human pathogens *Haemophilus* and *Pasteurella* (**Fig. 4a, Data S3**). Overall, we found that more than 10% of the analyzed bacteria have lost at least one ancestral hibernation factor gene (either HPF or RMF) (**Fig. 4b-d**). Thus, we found that the degeneration of hibernation factors can lead to their complete loss in a substantial number of bacteria, implying that many bacterial species rely on alternative, yet unknown, mechanisms for ribosome storage.

**Figure 4.**
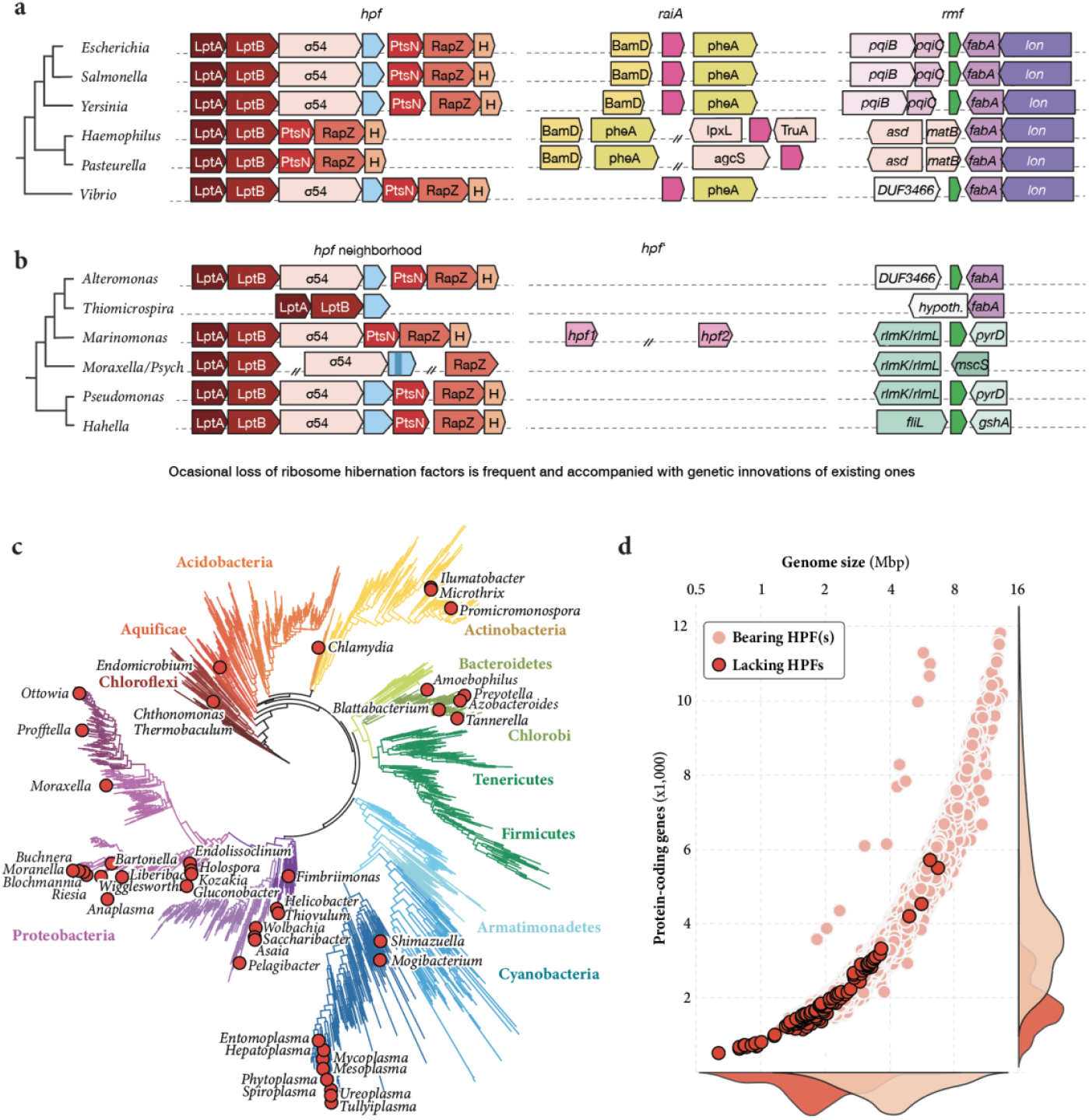
Ancestral hibernation factors have undergone extinction in a substantial fraction of bacteria. **(a, b)** Individual branches of γ-proteobacteria illustrate accidental loss of the ancestral *hpf* gene in some lineages of bacteria. (**a**) In *Haemophilus* and *Pasteurella* species, the ancestral *hpf* gene was lost along with its genomic neighbor, which encodes the stress-response protein RNA polymerase sigma factor 54 (σ54). This gene loss was accompanied by the genomic relocation of the *raiA* gene to a new genetic environment. (**b**) In *Marinomonas* and *Moraxella/Psychrobacter* species, the loss of the ancestral *rmf* gene was accompanied by the genomic transposition of the *hpf* gene. (**c**) The bacterial tree of life highlights lineages lacking HPF homologs, illustrating their widespread distribution across bacterial branches. (**d**) Comparison of genome size of organisms bearing and lacking HPF homologs illustrates a commonly occurring loss of HPF in the vast majority of organisms with small genomes (among other bacteria).

### Bacteria appear to buffer truncations in hibernation factors by multiplying their gene copy numbers

We then asked whether the reductive evolution of a particular hibernation factor impacts the evolution of the remaining hibernation machinery. We reasoned that if laboratory-induced knockouts of hibernation factor genes result in a loss of viability in cell cultures, then a similar loss occurring in a natural environment should also result in a loss of viability—unless cells evolve compensatory mechanisms for ribosome hibernation.

Our analysis revealed that when one or a few factors undergo truncations or gene loss, it is typically accompanied by substantial changes in the remaining machinery. We found that when bacterial species acquire a truncation in a hibernation factor (e.g., the C-terminal domain of HPF), they tend to obtain additional gene copies of this factor via horizontal gene transfer. For instance, we found that nearly one-third of bacterial species possess multiple *hpf* gene copies, with one copy corresponding to the ancestral gene (e.g., HPF in *E. coli*) and additional copies corresponding to the genes are acquired either via horizontal transfer or ancient gene duplication event (e.g., RaiA in *E. coli*; **Data S1**). Our phylogenetic mapping revealed that the truncation of the ancestral *hpf* is typically accompanied by the acquisition of additional *hpf* gene copies. This trend can be illustrated by comparing major bacterial clades. In Terrabacteria, where HPF is predominantly full-length, multiple *hpf* gene copies are found in fewer than 0.2% of species analyzed. But within Gracilicutes, which frequently possess truncated HPF, multiple *hpf* copies are predominant in each phylum bearing truncated HPF (**Fig. 5a-d**). Thus, we found that bacteria appear to buffer HPF truncations by multiplying its gene copy numbers, possibly to insure higher intracellular levels of this protein so that it remains bound to the ribosome even in the absence of the C-terminal domain.

**Figure 5.**
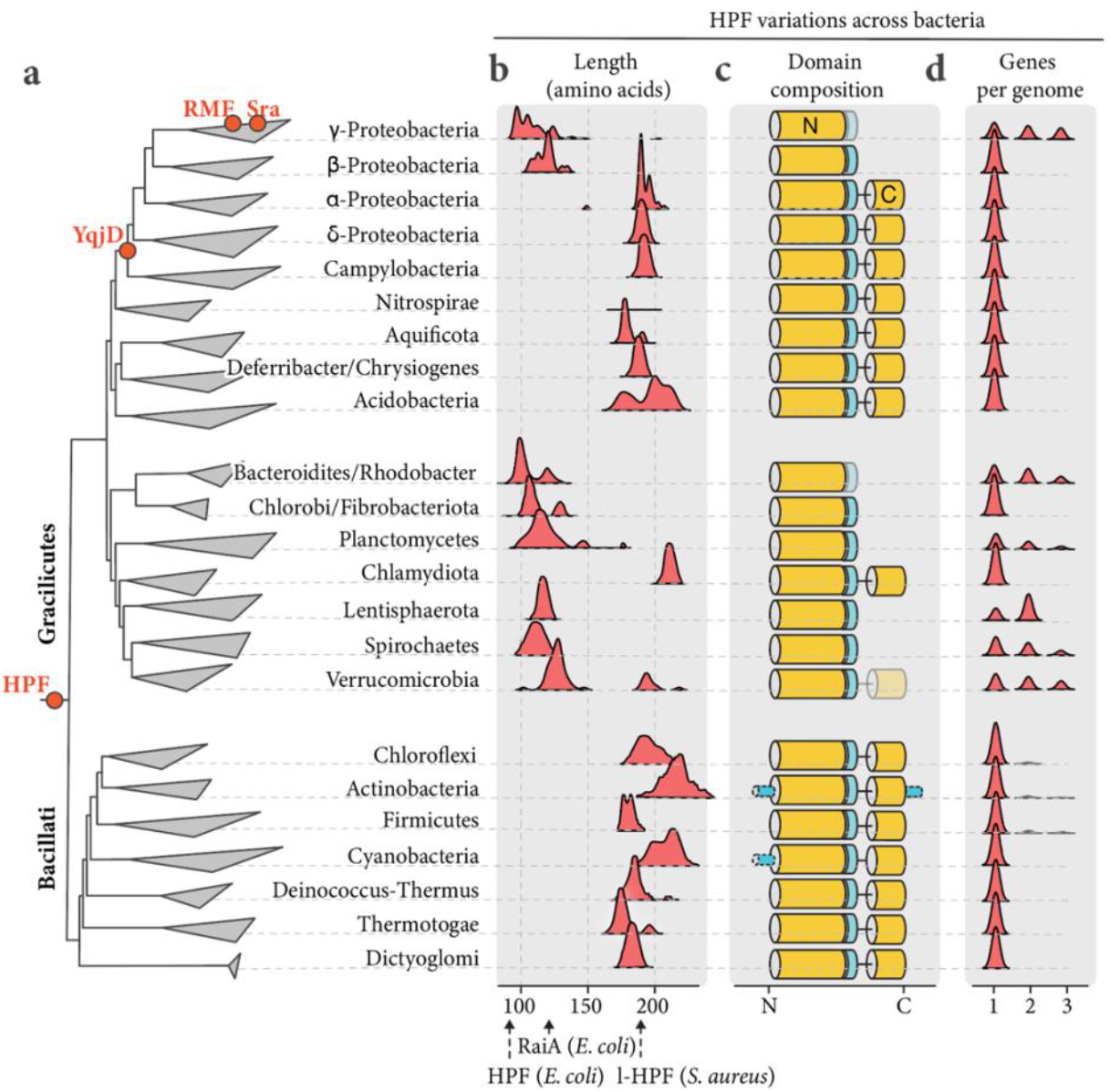
Bacteria appear to buffer reductive evolution of hibernation factors by multiplying their copy gene number. (**a**) The bacterial tree illustrating the evolutionary events indicating the origin of ribosome hibernation factors in bacterial species. This panel shows that the ribosome hibernation machinery has gradually emerged over a period of nearly four billion years of bacterial evolution (**c, d**) The schematics showing a typical length and domain structure of the ancestral HPF across bacterial phyla. (**d**) Plots summarizing the distributions of bacterial species within each phylum by the number of HPF-coding genes per genome. Overall, the panels **c-e** reveal that the hibernation factor HPF undergoes reductive evolution in several bacterial phyla, and this reductive evolution is often accompanied by the acquisition of additional HPF gene copies (e.g. in γ-proteobacteria, Bacteroidetes, Planctomycetes or Lentisphaerae). In other lineages, truncated HPF (e.g. β-proteobacteria or Chlorobi) acquires additional loop that likely attaches ribosomes to hibernation factor.

### Bacteria can restore truncations in long HPF by fusing it with other stress-response proteins

We found that some homologs of long HPF form stand-alone branches with sequences highly divergent from the major HPF groups. Our examination of their domain composition revealed that they represent previously uncharacterized HPF variants (**Fig. 6a, Fig. S2**). Although most these proteins are now erroneously annotated as long (full-size) HPF, these divergent HPF isoforms in fact lack the C-terminal domain responsible for ribosome dimerization and are instead fused to domains from stress-response proteins, such as the cold-shock protein Csp, the stress-response protein SigE, or CBS domains recently found in the hibernation factors Hib and Dri in archaea.^30,32^ Thus, we found that long HPF—now viewed as a structurally conserved bacterial hibernation factor—is in fact a family of structurally dissimilar HPF variants.

**Figure 6.**
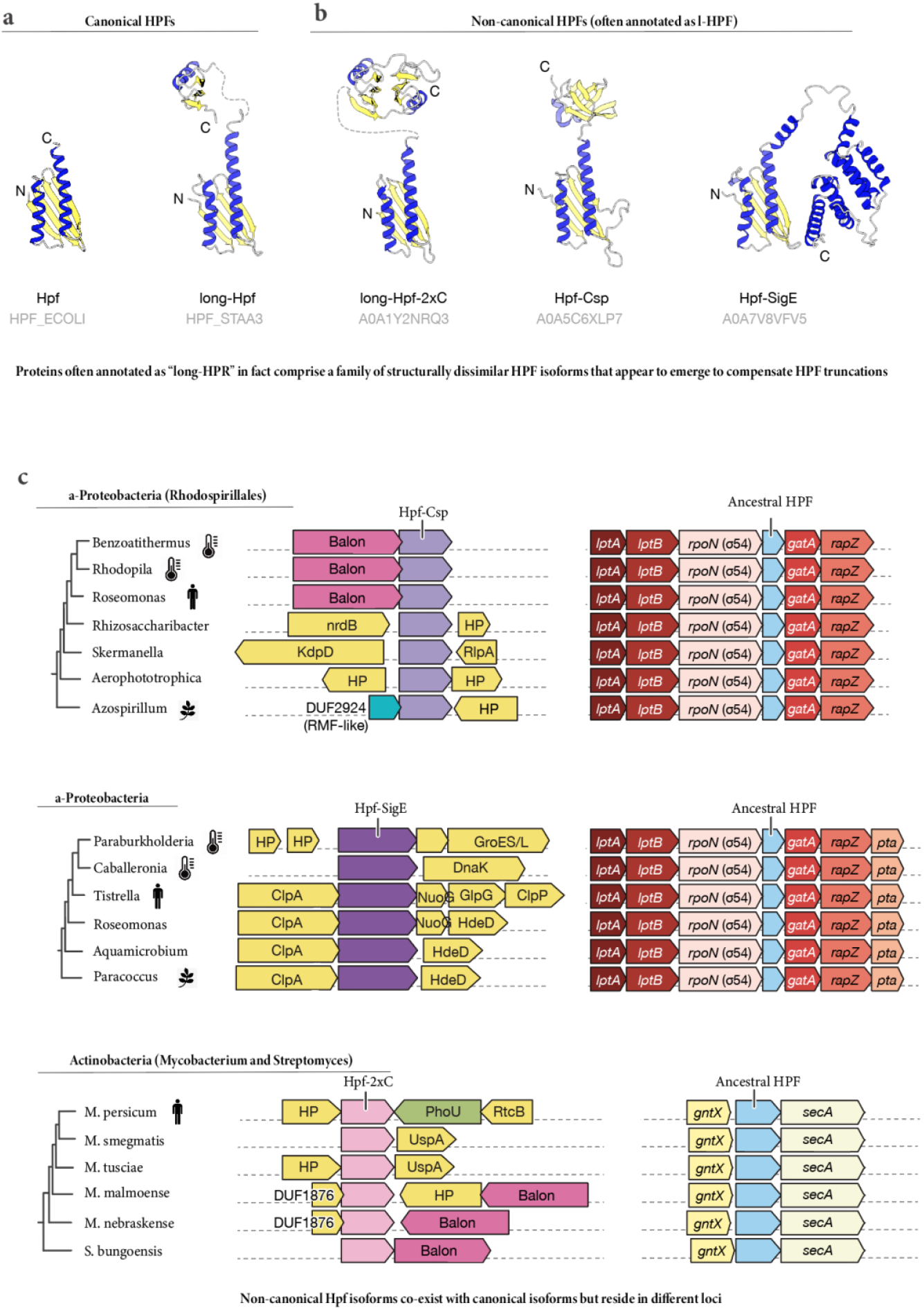
The seemingly conserved protein HPF is in fact a family of structurally dissimilar proteins. (**a, b**) Protein structures show canonical (**a**) and non-canonical (**b**) isoforms of hibernation factor HPF. (**c**) A few representative operon structures compare the genomic location of genes for non-canonical and canonical hibernation factors HPF.

Our phylogenetic analysis revealed that these chimeric HPF isoforms are scattered across bacterial species, including human pathogens, suggesting their frequent propagation primarily via horizontal gene transfers (**Fig. 6b,c**). Consistent with this idea, our analysis revealed that chimeric HPF isoforms can not only be found aside in genomes, but also on plasmids, including integrative plasmids, thereby explaining how these unusual HPF isoforms can spread across bacterial genomes. One example of such plasmid is pEc78, which confers resistance to antibiotics and environmental stress in pathogenic strains of *E. coli*.^38^ We found that pEc78 encodes a SigE-HPF fusion (**Fig. 6a**). This finding shows that, in addition to inventing new hibernation factors in certain lineages, bacteria can spread the genes for these factors via previously unappreciated routes of plasmid exchange. This has led to unexpected similarities in the ribosome hibernation apparatus between distant bacterial species and has made it possible for bacterial species to rapidly acquire new hibernation factors both in nature and clinical settings.

Importantly, within all bacterial genomes analyzed, chimeric HPF isoforms do not replace canonical HPF isoforms (descending from ancestral HPF) but coexist with them—so that a single species contains a canonical HPF and an additional HPF chimera (**Fig. 3b**). However, the genes for chimeric HPF reside on different genomic loci: canonical HPF homologs (typically, long HPF) retain their location in conserved *hpf* operons, while the chimeric *hpf* genes have a variable genomic location and are frequently found adjacent to genes for other non-canonical hibernation factors, such as Balon. Furthermore, our re-examination of RNA atlas studies of human pathogens^16^ revealed that bacteria can activate their genes for HPF isoforms in a reciprocal and condition-specific manner. For example, *Mycobacterium tuberculosis* possesses two HPF isoforms, including the canonical long HPF (also known as Rv3241c) and an HPF isoform which bears a duplicated C-terminal domain (also known as S30AE-containing protein, RafH, Datin and Rv0079^22,39,40^). Both these genes are activated by stationary phase, consistent with their role in ribosome hibernation. However, under hypoxia—the condition that most commonly causes *M. tuberculosis* to enter dormant, non-replicative states in the human body—the expression of canonical HPF is strongly inhibited, while the non-canonical isoform is activated. This reciprocal regulation suggests that bacterial cells not only can invent new hibernation factors by fusing HPF to stress-response domains but also utilize these non-canonical HPF isoforms to initiate stress-specific modes of ribosome hibernation.

### In studied species, the loss of ancestral hibernation factors appears to force bacteria to acquire novel ones

Although most species that have lost ancestral hibernation factors have not been experimentally assessed to understand how they cope without these protective proteins, we could derive insight from the recently characterized clade of Moraxellaceae. Members of this family of proteobacteria, particularly the genus *Psychrobacter*, were shown to possess an atypical hibernation factor, Balon, which is distinct from those of most other proteobacteria^14^. We used this clade to investigate the sequence of evolutionary events that led to the origin of Balon.

Our analysis revealed that approximately 70-100 Mya, the common ancestor of Moraxellaceae lost the gene for the hibernation factor RMF, which is now absent in all the members of this family and distinguishes this family from its relatives on the tree of life (**Fig. 7**). This loss was accompanied by disintegration of the HPF-coding operon (which became separated into three gene clusters scattered through bacterial genomes), and a modification of the HPF structure: this protein acquired an additional protein loop anchoring HPF to the decoding site of the ribosome^14^ and possibly compensating for the lack of RMF (**Fig .7**). Shortly after (∼65-69 Mya), when Moraxellaceae divided into the species of *Acinetobacter* and *Moraxella/Psychrobacter*, the ancestor of *Moraxella/Psychrobacter* acquired the gene for the hibernation factor Balon (likely via horizontal gene transfer from Actinobacteria due to the sequence similarity of Balon between these taxa), while some of the *Acinetobacter* species acquired the gene for aRDF, a protein recently characterized as a ribosome hibernation factor from Archaea^30^. Thus, we found that the loss of ancestral hibernation factors (e.g., RMF) appears to lead to a substantial remodeling of the hibernation machinery, forcing species to acquire novel hibernation factors and leading to a greater diversity of hibernation mechanisms even between closely related species.

**Figure 7.**
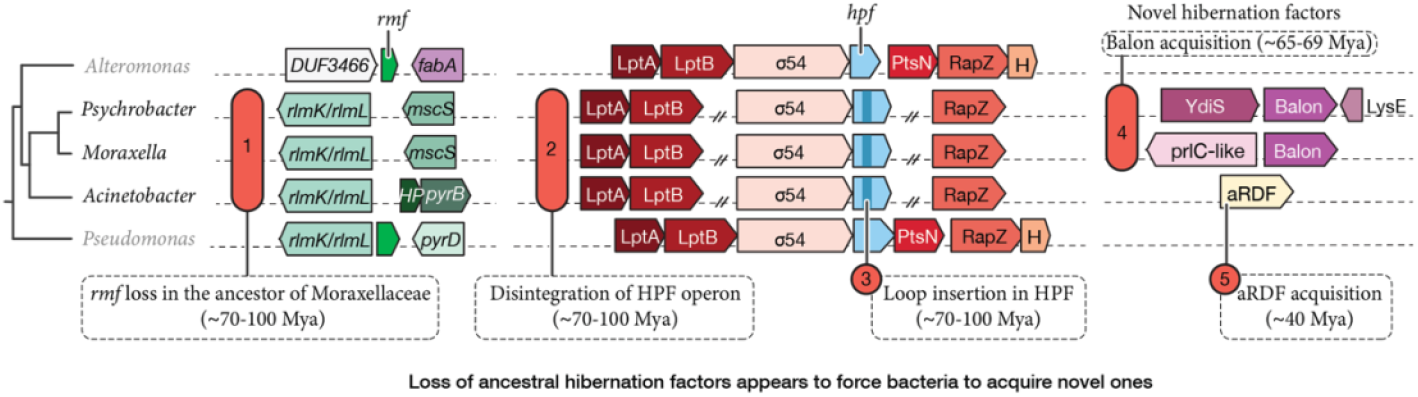
The loss of ancestral hibernation factors appears to force bacteria to acquire novel ones, creating mechanistic diversity of ribosome hibernation in closely related species. The bacterial tree and schematic operon structures illustrate the origin of the hibernation factor Balon and the loss of ancestral factor RMF in the family of Moraxellaceae (includes *Acinetobacter, Moraxella* and *Psychrobacter* species).

This example is important because it illustrates that hibernation factors evolve not in a random but a highly coordinated manner, where a change in one factor appears to trigger changes in the remaining hibernation machinery. Furthermore, this example of Moraxellaceae explains why some bacteria possess “unusual” hibernation factors compared to the more conventional ones found in their close relatives on the tree of life. It implies that similar changes (i.e. acquisitions of novel hibernation factors) could have occurred in other bacteria that have lost their ancestral hibernation factors but currently remain unstudied in terms of ribosome hibernation mechanisms.

## METHODS

### Bacterial growth and fitness measurements

*E. coli* strains, including the wild-type (K-12 strain) and strains with deletions of *raiA, hpf, rmf*, and *yqjD* (obtained from^41^), were inoculated in LB medium at 37 °C with shaking at 200 rpm and allowed to reach the stationary phase. The obtained stationary cultures were then continuously maintained at 37 °C with shaking 200 rpm, and aliquots were taken periodically to measure viability and recovery times for each strain. For each recovery time measurement, the aliquots of cell cultures were transferred into a 24-well plate, with each well containing 10 µL of a stationary-phase culture mixed with 1.25 mL of LB medium. The plates were then incubated in a plate reader, and the OD_630_ was continuously monitored to assess bacterial growth. To achieve better temporal separation, the growth of the strains was measured at room temperature to slow bacterial growth. In each experiment, each strain was analyzed in four different wells of a 24 well plate.

### Initial search for and analysis of hibernation factor homologs

To identify homologs of hibernation factors, we used a compinged output of two alternative approaches. In the first approach, we used HMMER^42^ to analyze protein sequences deposited to the database TrEMBL.^43^ The following settings were used for each search: algorithm: phmmer, parameters: -E 1 --domE 1 --incE 0.01 -- incdomE 0.03 --mx BLOSUM62 --pextend 0.4 --popen 0.02 --seqdb referenceproteomes. We used JACKHMMER which completes up to 3 iterations to generate proteins based on the query sequence + homologs found from the previous iterations. We then separated the accession codes and accessed them on Uniprot ID mapping to generate excel tables with information including the species name and phyla. To separate homologs of vertically evolving clusters, we used multiple sequence alignments of protein sequences with ClustalO. To estimate the time points of when bacteria have acquired, truncated, or lost their ribosome hibernation factors, we used the recently published phylogenetic trees assessing the age of species.^44,36^

### Analysis of sequences and domain composition of HPF/RaiA homologs

The Genome Taxonomy Database (GTDB) Release 09-RS220 ^45^ was used to collect proteomes from high-quality representative genomes that met the following criteria: completeness ≥95%, contamination ≤5%, and the presence of the 16S rRNA genes and ≥18 tRNAs (genomes are listed in **Data S1**). We then used the hidden Markov model profile of PF02482 from Pfam as a query in hmmsearch (HMMER 3.4) to find HPF/RaiA candidates from the proteomes.^46,47^ We then used InterProScan 5.73-104.0 for further screening ^48^, identifying hits to IPR036567 as HPF/RaiA homologs. The HPF/RaiA homologs were classified according to the domain architectures revealed by InterProScan. Data analysis and visualization were performed using R 4.3.2 ^49^ with the following packages: tidyverse 2.0.0 W ^50^, plyranges 1.22.0^51^, and ggupset 0.3.0^52^. Species trees (bac120.tree and ar53.tree) were downloaded from GTDB Release 09-RS220. The trees were pruned and visualized using the Python package DendroPy 5.0.6^53^ and the R packages ape 5.8^54^, ggtree 3.10.1 ^55^, ggtreeExtra 1.12.0^56^, tidytree 0.4.6 ^57^, and ggnewscale 0.4.10^58^.

### Analysis of HF-encoding operons

The operon structure, composition and evolution was carried out using WebFlags^59^ using the following input parameters: blastp database: refseq_select (representative records), E-value cutoff for Blastp searching: 1e-3, Jackhmmer E-value: 1e-10, number of Jackhmmer iterations: 3, Number of flanking genes: 3-5.

### dN/dS evaluation

The dN/dS ratios for the sequences of hibernation and translation factors were estimated using custom python scripts using the original definition by Kimura.^60,61^ The input sequences were retrieved using *datasets* and *dataformat* commands in NCBI Datasets CLI of the NCBI Datasets API.^62^

### Proteomic analysis of ribosomes in stationary *E. coli*

An overnight culture of *E. coli* cells (3 OD_600_) was washed in buffer A (50 mM Tris-HCl pH 7.5, 20 mM magnesium acetate and 50 mM KCl) and was transferred to 2 mL microcentrifuge tubes containing approximately 0.1 mL of 0.5 mm zirconium beads (Sigma-Aldrich BeadBug). The cells were disrupted by shaking for 30 s at 6.5 ms^-1^ speed in a bead beater (Thermo FastPrep FP120 Cell Disrupter). The cells were then centrifuged for 5 min at 16,000 x g and 4 °C to remove cell debris. The resulting supernatant was then centrifuged again for 1 min at 16,000 x g and 4 °C to remove the remaining debris. We then analysed 0.25 mL crude *E. coli* lysates on top of 12 mL 10-50% sucrose gradients in buffer A and were centrifuged for 18 h at 14,200 rpm at 4 °C. The sucrose gradients were fractionated in 0.5 mL increments and the absorbance of each fraction was measured at OD_260_. The samples corresponding to ribosome peaks were pooled together, concentrated using Amicon Ultra 4 centrifugal filters by centrifuging for 30 m at 8,000 x g and aliquoted into 10 µL fractions, flash frozen with liquid nitrogen for proteomic analysis.

Protein isolation for mass-spectrometry was carried out using S-Trap micro spin columns (Protifi). SDS was added to the samples to a final concentration of 5%, and volume adjusted to 20 µl. 20 µl of S-trap lysis buffer (50mM TEAB, 5% SDS) was added to the beads. From that point the same treatment was applied to all samples. Proteins were reduced with dithiothreitol (DTT) at final concentration of 20 mM (65°C, 30 min). Cysteines were alkylated by incubation with iodoacetamide (40 mM final concentration, 30 min, room temp. in dark) and then acidified by adding 27.5% Phosphoric acid to a final concentration of 2.5% (v/v). The samples were then loaded onto spin columns in 6 volumes of loading buffer (90% methanol 100mM TEAB pH 8) and centrifuged at 4,000 x g for 30 s. The columns were then washed with loading buffer (three times) and the flow through discarded. Proteins were digested with trypsin (Worthington) in 50mM TEAB pH 8.5, at a ratio of 10:1 protein to trypsin, overnight at 37°C.

Peptides were eluted with three washes of the trap; first 50 µl 50mM TEAB, second 50ul 0.1% formic acid and third 50 µl 50% acetonitrile with 0.1% formic acid. The solution was frozen, then dried in a centrifugal concentrator and reconstituted in 20 µl of 0.1% formic acid 2% Acetonitrile. Then, 1 µl of each peptide sample was loaded per LCMS run. Peptides were separated using an UltiMate 3000 RSLCnano HPLC. Samples were first loaded/desalted onto in house made C8 trap column (15mm/0.275mm i.d., Dr. Maisch ReproSil-Pur 120 C8, 5 µm) at a flow rate of 10 μL min^−1^ maintained at 45 °C and then separated on a 75 μm x 75 cm C18 column (Thermo EasySpray -C18 2 µm) with integrated emitter using a 60 min nonlinear gradient from 95 % A (0.1% FA) and 5% B (0.1% FA in 80% ACN), to 35% B, at a flow rate of 150 nL min^−1^. The eluent was directed to an Thermo Q Exactive HF mass spectrometer through the EasySpray source at a temperature of 280 °C, spray voltage 1500 V. The total LCMS run time was 120 min. Orbitrap full scan resolution was 120,000, ACG Target 5e6, maximum injection time 100 ms, scan range 390-1050 m/z. DIA MS/MS were acquired with 15 m/z windows covering 319.5.5-1024.5 m/z, at 30000 resolution, maximum injection time of 100 ms, with ACG target set to 3e6, and normalized collision energy level of 27.

The acquired data has been analysed in DIA-NN version 2.1.0 (DIA-NN: neural networks and interference correction enable deep proteome coverage in high throughput Nature Methods, 2020) against Clostridium perfringens proteome sequence database (Uniprot UP000000625, version from 2025/03/26) combined with common Repository of Adventitious Proteins (cRAP), Fragment m/z: 300-1800, enzyme: Trypsin, allowed missed-cleavages: 1, peptide length: 7-30, precursor m/z 300-1250, precursor charge: 2-4, Fixed modifications: carbamidomethylation (C), Variable modifications: Oxidation (M), Acetylation (N-term). The processed data are shown in (**Data S2**).

## ACKNOWLEDGEMENTS

We thank Gabriel Demo and Ahmed Hassan (both from Central European Institute of Technology, Brno, Czech Republic) for sharing their analysis of aRDF homologs in bacterial species. This work was funded by the Biotechnology and Biological Sciences Research Council (BB/T008695/1 to L.I.C. and C.R.B.), the MRC Discovery Medicine North (DiMeN) Doctoral Training Partnership (MR/N013840/1 to C.L.E.), JST CREST Grant Number JPMJCR20S4 (K.O., K.A. and S.S.) and the International Collaboration Award from the Royal Society, UK (ICA/R2/242249 to S.S. and S.V.M.).

## DATA AVAILABILITY STATEMENT

The **Data S1-S3** are available on the FigShare repository: 10.6084/m9.figshare.30521297.

## REFERENCES

1. Shore, D. & Albert, B. Ribosome biogenesis and the cellular energy economy. Current Biology 32, R611– R617 (2022).

2. Burton, R. F. Biology by Numbers: An Encouragement to Quantitative Thinking. (Cambridge University Press, Cambridge, 1998). doi:10.1017/CBO9780511802713.

3. Wada, A., Yamazaki, Y., Fujita, N. & Ishihama, A. Structure and probable genetic location of a ‘ribosome modulation factor’ associated with 100S ribosomes in stationary-phase Escherichia coli cells. Proceedings of the National Academy of Sciences 87, 2657–2661 (1990).

4. Agafonov, D. E., Kolb, V. A., Nazimov, I. V. & Spirin, A. S. A protein residing at the subunit interface of the bacterial ribosome. Proceedings of the National Academy of Sciences 96, 12345–12349 (1999).

5. Ben-Shem, A. et al. The structure of the eukaryotic ribosome at 3.0 Å resolution. Science 334, 1524–1529 (2011).

6. Anger, A. M. et al. Structures of the human and Drosophila 80S ribosome. Nature 497, 80–85 (2013).

7. Brown, A., Baird, M. R., Yip, M. C., Murray, J. & Shao, S. Structures of translationally inactive mammalian ribosomes. eLife 7, e40486 (2018).

8. Beckert, B. et al. Structure of a hibernating 100S ribosome reveals an inactive conformation of the ribosomal protein S1. Nature microbiology 3, 1115–1121 (2018).

9. Barandun, J., Hunziker, M., Vossbrinck, C. R. & Klinge, S. Evolutionary compaction and adaptation visualized by the structure of the dormant microsporidian ribosome. Nat Microbiol 4, 1798–1804 (2019).

10. Ehrenbolger, K. et al. Differences in structure and hibernation mechanism highlight diversification of the microsporidian ribosome. PLOS Biology 18, e3000958 (2020).

11. Nicholson, D. et al. Adaptation to genome decay in the structure of the smallest eukaryotic ribosome. Nat Commun 13, 591 (2022).

12. Leesch, F. et al. A molecular network of conserved factors keeps ribosomes dormant in the egg. Nature 613, 712–720 (2023).

13. McLaren, M. et al. CryoEM reveals that ribosomes in microsporidian spores are locked in a dimeric hibernating state. Nat Microbiol 8, 1834–1845 (2023).

14. Helena-Bueno, K. et al. A new family of bacterial ribosome hibernation factors. Nature 626, 1125–1132 (2024).

15. Blandy, A. et al. Translational activity of 80S monosomes varies dramatically across different tissues. 2024.06.24.600330 Preprint at 10.1101/2024.06.24.600330 (2024).

16. Avican, K. et al. RNA atlas of human bacterial pathogens uncovers stress dynamics linked to infection. Nature communications 12, 3282 (2021).

17. A, S. et al. The quantitative and condition-dependent Escherichia coli proteome. Nature biotechnology 34, (2016).

18. Helena-Bueno, K., Chan, L. & Melnikov, S. Rippling life on a dormant planet: hibernation of ribosomes, RNA polymerases and other essential enzymes. Frontiers in Microbiology 15, 1386179 (2024).

19. Wada, A., Igarashi, K., Yoshimura, S., Aimoto, S. & Ishihama, A. Ribosome Modulation Factor: Stationary Growth Phase-Specific Inhibitor of Ribosome Functions from Escherichia coli. Biochemical and Biophysical Research Communications 214, 410–417 (1995).

20. El-Sharoud, W. M. & Niven, G. W. The influence of ribosome modulation factor on the survival of stationary-phase Escherichia coli during acid stress. Microbiology 153, 247–253 (2007).

21. Prossliner, T., Gerdes, K., Sørensen, M. A. & Winther, K. S. Hibernation factors directly block ribonucleases from entering the ribosome in response to starvation. Nucleic Acids Research 49, 2226–2239 (2021).

22. Trauner, A., Lougheed, K. E., Bennett, M. H., Hingley-Wilson, S. M. & Williams, H. D. The dormancy regulator DosR controls ribosome stability in hypoxic mycobacteria. Journal of Biological Chemistry 287, 24053–24063 (2012).

23. Basu, A. & Yap, M.-N. F. Disassembly of the Staphylococcus aureus hibernating 100S ribosome by an evolutionarily conserved GTPase. Proceedings of the National Academy of Sciences 114, E8165–E8173 (2017).

24. Puri, P. et al. Lactococcus lactis YfiA is necessary and sufficient for ribosome dimerization. Molecular Microbiology 91, 394–407 (2014).

25. Basu, A., Shields, K. E., Eickhoff, C. S., Hoft, D. F. & Yap, M.-N. F. Thermal and Nutritional Regulation of Ribosome Hibernation in Staphylococcus aureus. Journal of Bacteriology 200, 10.1128/jb.00426-18 (2018).

26. Flygaard, R. K., Boegholm, N., Yusupov, M. & Jenner, L. B. Cryo-EM structure of the hibernating Thermus thermophilus 100S ribosome reveals a protein-mediated dimerization mechanism. Nat Commun 9, 4179 (2018).

27. Li, Y. et al. Zinc depletion induces ribosome hibernation in mycobacteria. Proc Natl Acad Sci U S A 115, 8191–8196 (2018).

28. Feaga, H. A., Kopylov, M., Kim, J. K., Jovanovic, M. & Dworkin, J. Ribosome Dimerization Protects the Small Subunit. Journal of Bacteriology 202, 10.1128/jb.00009-20 (2020).

29. Majumdar, S. et al. A Novel Actinobacteria-Specific Ribosome Hibernation Factor in Mycobacterium tuberculosis. bioRxiv 2024–11 (2024).

30. Hassan, A. H. et al. Novel archaeal ribosome dimerization factor facilitating unique 30S–30S dimerization. Nucleic Acids Research 53, gkae1324 (2025).

31. Nissley, A. J. et al. Structure of an archaeal ribosome reveals a divergent active site and hibernation factor. Nature Microbiology 10, 1940–1953 (2025).

32. madru, C. et al. A new family of ribosome hibernation factors in Archaea. bioRxiv 2025–10 (2025).

33. Ueta, M. et al. Conservation of two distinct types of 100S ribosome in bacteria. Genes to Cells 18, 554–574 (2013).

34. Prossliner, T., Skovbo Winther, K., Sørensen, M. A. & Gerdes, K. Ribosome Hibernation. Annual Review of Genetics 52, 321–348 (2018).

35. Njenga, R. K. et al. Ribosome-inactivation by a class of widely distributed C-tail anchored membrane proteins. Structure https://www.cell.com/structure/fulltext/S0969-2126(24)00388-5 (2024).

36. Davín, A. A. et al. A geological timescale for bacterial evolution and oxygen adaptation. Science 388, eadp1853 (2025).

37. Yamagishi, M. et al. Regulation of the Escherichia coli rmf gene encoding the ribosome modulation factor: growth phase- and growth rate-dependent control. The EMBO Journal 12, 625–630 (1993).

38. Papagiannitsis, C. C., Kutilova, I., Medvecky, M., Hrabak, J. & Dolejska, M. Characterization of the Complete Nucleotide Sequences of IncA/C_2_ Plasmids Carrying In809-Like Integrons from Enterobacteriaceae Isolates of Wildlife Origin. Antimicrob Agents Chemother 61, e01093–17 (2017).

39. A, K. et al. Mycobacterium tuberculosis DosR regulon gene Rv0079 encodes a putative, ‘dormancy associated translation inhibitor (DATIN)’. PloS one 7, (2012).

40. Kumar, N., Sharma, S. & Kaushal, P. S. Cryo-EM structure of the mycobacterial 70S ribosome in complex with ribosome hibernation promotion factor RafH. Nature Communications 15, 638 (2024).

41. Baba, T. et al. Construction of Escherichia coli K-12 in-frame, single-gene knockout mutants: the Keio collection. Molecular Systems Biology 2, 2006.0008 (2006).

42. Potter, S. C. et al. HMMER web server: 2018 update. Nucleic acids research 46, W200–W204 (2018).

43. O’Donovan, C. et al. High-quality protein knowledge resource: SWISS-PROT and TrEMBL. Briefings in bioinformatics 3, 275–284 (2002).

44. Strassert, J. F., Irisarri, I., Williams, T. A. & Burki, F. A molecular timescale for eukaryote evolution with implications for the origin of red algal-derived plastids. Nature Communications 12, 1879 (2021).

45. Parks, D. H. et al. A standardized bacterial taxonomy based on genome phylogeny substantially revises the tree of life. Nature biotechnology 36, 996–1004 (2018).

46. Finn, R. D. et al. Pfam: the protein families database. Nucleic acids research 42, D222–D230 (2014).

47. Eddy, S. R. Accelerated profile HMM searches. PLoS computational biology 7, e1002195 (2011).

48. Jones, P. et al. InterProScan 5: genome-scale protein function classification. Bioinformatics 30, 1236–1240 (2014).

49. Team, R. C. R: A language and environment for statistical computing. R Foundation for Statistical Computing, Vienna, Austria. http://www.R-project.org/ https://cir.nii.ac.jp/crid/1574231874043578752 (2016).

50. Wickham, H. et al. Welcome to the Tidyverse. JOSS 4, 1686 (2019).

51. Lee, S., Cook, D. & Lawrence, M. plyranges: a grammar of genomic data transformation. Genome Biol 20, 4 (2019).

52. Ahlmann-Eltze, C. ggupset. Combination matrix axis for ‘ggplot2’to create’UpSet’plots. R package version 0.3. 0. (2020).

53. Moreno, M. A., Holder, M. T. & Sukumaran, J. DendroPy 5: a mature Python library for phylogenetic computing. JOSS 9, 6943 (2024).

54. Paradis, E. & Schliep, K. ape 5.0: an environment for modern phylogenetics and evolutionary analyses in R. Bioinformatics 35, 526–528 (2019).

55. Yu, G., Smith, D. K., Zhu, H., Guan, Y. & Lam, T. T. GGTREE : an R package for visualization and annotation of phylogenetic trees with their covariates and other associated data. Methods Ecol Evol 8, 28–36 (2017).

56. Xu, S. et al. ggtreeExtra: compact visualization of richly annotated phylogenetic data. Molecular biology and evolution 38, 4039–4042 (2021).

57. Yu, G. Data Integration, Manipulation and Visualization of Phylogenetic Trees. (Chapman and Hall/CRC, 2022).

58. Campitelli, E. ggnewscale: Multiple Fill and Colour Scales in’ggplot2’. R package version 0.4 5, (2020).

59. Saha, C. K., Sanches Pires, R., Brolin, H., Delannoy, M. & Atkinson, G. C. FlaGs and webFlaGs: discovering novel biology through the analysis of gene neighbourhood conservation. Bioinformatics 37, 1312–1314 (2021).

60. Kimura, M. Preponderance of synonymous changes as evidence for the neutral theory of molecular evolution. Nature 267, 275–276 (1977).

61. Jeffares, D. C., Tomiczek, B., Sojo, V. & Dos Reis, M. A Beginners Guide to Estimating the Non-synonymous to Synonymous Rate Ratio of all Protein-Coding Genes in a Genome. in Parasite Genomics Protocols (ed. Peacock, C.) vol. 1201 65–90 (Springer New York, New York, NY, 2015).

62. O’Leary, N. A. et al. Exploring and retrieving sequence and metadata for species across the tree of life with NCBI Datasets. Scientific data 11, 732 (2024).

